# Pattern-centric transformation of omics-data sources grounded on multi-wise gene associations aids predictive tasks in TCGA while ensuring interpretability

**DOI:** 10.1101/2023.05.28.542574

**Authors:** André Patrício, Rafael S. Costa, Rui Henriques

**Affiliations:** INESC-ID, Instituto Superior Técnico, Universidade de Lisboa, R. Alves Redol 9, 1000-029, Lisboa, Portugal; LAQV-REQUIMTE, Department of Chemistry, NOVA School of Science and Technology, NOVA University Lisbon, Campus da Caparica, 2829-516, Caparica, Portugal

**Keywords:** omics-data, TCGA, dimensionality reduction, interpretability, systems biology, cancer, data integration, predictive models

## Abstract

**Motivation:** The increasing prevalence of omics data sources is pushing the study of regulatory mechanisms underlying complex diseases such as cancer. However, the vast quantities of features produced and the inherent interplay between them lead to a level of complexity that hampers both descriptive and predictive tasks, requiring custom-built algorithms that can extract relevant information from these sources of data.

**Results:** We propose a transformation that moves data centered on molecules (e.g. transcripts and proteins) to a new data space focused on putative regulatory modules given by statistically relevant patterns of coexpression. The proposed transformation extracts patterns from the data through biclustering and uses them to create new variables with guarantees of interpretability and discriminative power. The transformation is shown to achieve dimensionality reductions of up to 99% and to increase the predictive performance of various classifiers across multiple omics layers. Our results suggest that a transformation of omics data from gene-centric to pattern-centric data provides benefits to both prediction tasks and human interpretation. The proposed approach is expected to greatly support further bioinformatic analyses for precision medicine applications.

**Availability:** Software code and the raw results generated are available at github.com/Andrempp/Pattern-Centric-Transformation.

**Supplementary information:** Supplementary data are available at *Journal Name* online.

## 1. Introduction

Cancer is a class of diseases characterized by the abnormal growth, replication, and survivability of cells, which in 2020 presented 19,292,800 new cases and caused 9,958,100 deaths worldwide (Ferlay et al., 2021). This convoluted disease can have its origin in various tissues and arise due to disruptions in multiple cellular processes. New profiling techniques enable an in-depth look into the biological mechanisms related to complex diseases. The information generated by these techniques, termed omics data, can be categorized according to the targeted molecular structures, whether genes (genomics), transcripts (transcriptomics), or proteins (proteomics), amongst others. Data from these omics layers have been widely used in cancer research for multiple purposes, such as molecular subtyping (Zhao et al., 2019; Sun et al., 2022; Doebley et al., 2022), biomarker identification (Swan et al., 2013; Zhao et al., 2020; Ghosal et al., 2020), prognosis (Marquardt et al., 2012; Nault et al., 2013; Roessler et al., 2010), and drug development (Neary et al., 2021; Huang et al., 2019; Li et al., 2021). Still, the large quantities of data produced, hundreds or thousands of measurements per profiled sample, hinder the ability of humans and even algorithms to extract knowledge from it (Clarke et al., 2008).

Classic approaches to this problem focus on selecting Differentially Expressed Genes (DEGs) between the conditions under study (Zhang et al., 2015, 2018; Zhao et al., 2014). Although useful to reduce the high-dimensionality of omics data, these techniques are generally unable to capture dependencies among multiple genes. Since genes often work with each other in regulatory modules, this type of analysis can exclude genes that alone have a subtle differential profile, but when considered with other genes may have a decisive influence on one or more conditions, a problem shared by many other dimensionality reduction techniques. Procedures for reducing dimensionality focused on multivariate relationships (Ding and Peng, 2005; Gevaert et al., 2006; Yeung and Bumgarner, 2003) can be used to deal with this problem, but these are generally associated with the loss of potentially relevant information and interpretability (Saeys et al., 2007). Complementary advances, including integration of domain knowledge (Fan et al., 2019; Raghu et al., 2017; Yousef et al., 2021b), integration of multiple omics (Rohart et al., 2017; Metselaar et al., 2021), and ensemble of feature selection techniques (Lu et al., 2017; Metselaar et al., 2021), have been proposed, with some showing capability of improving predictive performance (Chandra and Gupta, 2011; Ding and Peng, 2005; Acharya et al., 2017; Yousef et al., 2021a).

This work provides a data transformation that can be applied to different types of omics data to increase the prediction capability of machine learning models while solving the previously mentioned challenges: dimensionality reduction; presence of putative regulatory modules with synergistic gene interplay on specific conditions; and preserved interpretability of the transformed data space.

We hypothesize that the defined goal can be achieved through pattern mining techniques, more precisely, biclustering. Biclustering algorithms search the data for subsets of observations following a coherent pattern across a subset of features, which in gene expression data corresponds to a group of co-expressed genes with a statistically significant regulatory pattern. The discriminative power of this pattern can then be calculated through various metrics. In accordance, biclustering has been widely used for putative regulatory module discovery (Madeira and Oliveira, 2004), with recent contributions improving the classic stances to further ensure functional relevance (Henriques and Madeira, 2014), statistical significance (Henriques and Madeira, 2018), and discriminative power (Henriques and Madeira, 2021).

The data transformation proposed in this work uses biclustering to extract relevant patterns from the data and subsequently use them to create new variables encoding the activation of said putative regulatory pattern. Contrary to other feature extraction methods, information loss is minimal as each pattern encompasses relevant information on a subset of the original molecular variables. Furthermore, interpretability is preserved in the reduced data space (arguably augmented given its organization in putative functions) and well suited for subsequent predictive ends as each pattern shows notable guarantees of discriminative power.

To validate the aforementioned qualities of the proposed pattern-centric data transformation, we study its impact on dimensionality reduction and classification performance across different predictive models. This analysis is performed with data from The Cancer Genome Atlas (TCGA) program (Weinstein et al., 2013), encompassing multiple omics layers from the Brain Lower Grade Glioma (TCGA-LGG) and Colon Adenocarcinoma (TCGA-COAD) projects.

## 2. Materials and Methods

The proposed data transformation converts a feature space based on molecules (genes, transcripts, proteins, etc..) to a space based on patterns of expression of these molecules. The transformation identifies relevant combinations of molecules and corresponding expression values and groups them into a single feature, resulting in new features that encapsulate information about patterns that are relevant to the problem at hand. The first step of the transformation is the mining of putative regulatory patterns in the data through pattern mining based biclustering (Henriques et al., 2015), followed by the creation of new features for each found pattern, where the corresponding value represents the presence of that regulatory pattern in the sample.

### 2.1. Datasets and Preprocessing

The data used for assessing the impact of the proposed transformation originates from The Cancer Genome Atlas (TCGA) program. Two distinct TCGA projects were selected based on heterogeneity and coverage of transcriptome- and proteome-wide data: Brain Lower Grade Glioma (TCGA-LGG)^1^ and Colon Adenocarcinoma (TCGA-COAD)^2^. For each project, we explore regulatory activity pertaining to miRNA, mRNA, and protein expression, and further access patients’ clinical variables. *Vital Status* is selected as the primary target in this work, encoding the survivability of the patient. The data was obtained through the R package TCGABiolinks (Colaprico et al., 2016) and corresponds to the Genomic Data Commons (GDC) harmonized database, which uses the human genome version GRCh38 as reference. The number of samples and variables per project and omics layer are shown in Table 1.

**Table 1.** Number of variables and samples in each dataset per project and omics layer.

| omics Layer | Nr. Variables | Samples per project |  |
| --- | --- | --- | --- |
|  |  | TCGA-LGG | TCGA-COAD |
| miRNA | 18881 | 530 | 465 |
| mRNA | 60664 | 534 | 524 |
| protein | 727<sup>1</sup> | 435 | 363 |

<sup>1</sup>487 in TCGA-LGG

From the available measurements in each layer, we select Reads Per Million Mapped Reads (RPM) for miRNA, Transcripts Per Kilobase Million (TPM) for mRNA, and relative levels of expression for proteins. Features with a high percentage of 0’s (≥95%) were removed, log transformation was applied to the miRNA and mRNA layers, and the filtering function from edgeR (Robinson et al., 2010; Chen et al., 2016) was used to perform preliminary filtering of genes not significantly expressed.

### 2.2. Biclustering

Given a multivariate dataset described by a set of observations *X* = {*x*_1_, …, *x*_*n*_} and a set of variables *Y* = {*y*_1_, …, *y*_*m*_}, biclustering is a data mining technique that identifies subspaces, each described by a subset of observations and variables *B* = (*I* ⊂ *X, J* ⊂ *Y*) with pattern *p* = {*y*_*j*_ = *c*_*j*_ |*y*_*j*_ ∈ *J* }, satisfying specific homogeneity and statistical significance criteria (e.g., coherently expressed genes on an unexpectedly high number of samples).

The homogeneity criteria of the patterns can be adjusted according to the addressed problem. Patterns can be described by constant values across observations or variables, shifting and scaling factors across rows or columns, and coherent order of variables. In this work, we use patterns of constant values across observations. Biclustering, therefore, has the advantage over clustering of only comparing observations according to a subset of variables. This characteristic is fundamental when considering gene expression data since genes work in subsets to form regulatory modules that control biological mechanisms.

Biclustering approaches based on pattern mining further show unique benefits, including efficient exhaustive searches, robustness to noise and missing values, flexible bicluster structure, and the possibility to customize the desirable pattern coherence, amongst others (Henriques et al., 2015). These characteristics make BicPAMS (Henriques et al., 2017), an algorithm that combines state-of-the-art contributions from pattern mining-based biclustering for biological data analysis, a viable candidate for the defined goal: the identification of relevant patterns of expression in omics data to transform gene-centric data spaces in pattern-centric data spaces.

### 2.3. Pattern-centric transformation

The proposed transformation works by extracting putative regulatory patterns from data and using them to generate new dimensions, moving from a gene-centric to a pattern-centric space. We start by applying the BicPAMS algorithm (Henriques et al., 2017) to identify patterns that yield statistical significance and are additionally discriminative of a selected target variable. Each pattern is defined by a set of observations, a set of variables and corresponding values, and descriptive metrics such as *p*-value and lift. The columns of the new dimension space will correspond to the found patterns. For each select pattern, the process followed is described in Figure 1: i) select a pattern and identify the corresponding values (Pattern 1 and values 3, 4, and 8); ii) create a column in the new dataset corresponding to this pattern (column *p1*); and iii) for each row, calculate the distance between its values and the values of the pattern (Euclidean distance between 6, 8, 5 and 3, 4, 8) and use it as the new value;

**Fig. 1:**
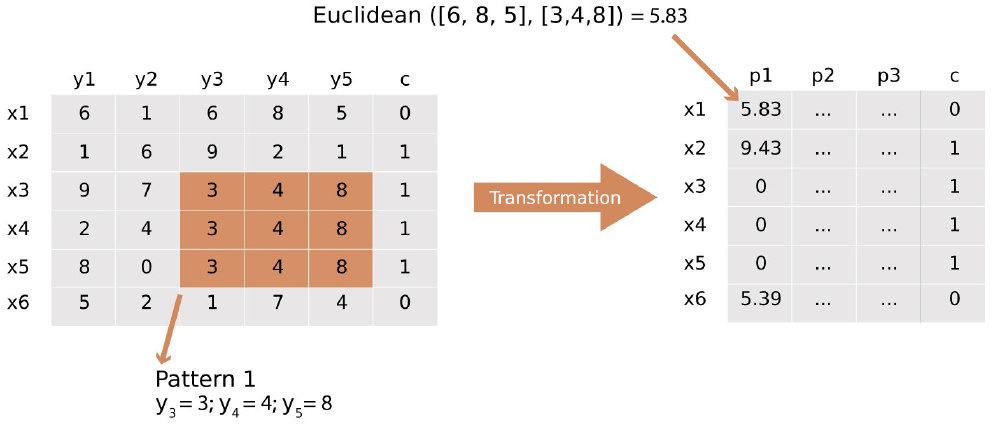
Example of pattern-centric transformation for a single pattern *p1*.

The transformation can be adjusted in various ways. Following its order of operations, it can start by filtering the patterns found by the biclustering algorithm based on their statistical significance, quality, discriminative power, or domain relevance (functional enrichment). For this purpose, we resort to the lift metric, which measures how well a pattern in the data predicts a given class. A pattern *p*, found to be related to a class *c*, would have a lift of,

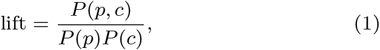

where *P* (*p*) is the probability of pattern *p* occurring in the data, *P* (*c*) is the probability of class *c* occurring in the data, and *P* (*p, c*) is the joint probability of *p* and *c* in the data.

In order to evaluate the expression level of a regulatory pattern in a sample, we further need to select a distance metric. According to sensitivity analysis over various similarity and correlation measures, the Euclidean distance is selected for this end. A final possible step is to perform a feature selection post-transformation, reducing the number of new features according to some criteria. However, in order to preserve the analysis of the proposed transformation, postprocessing options were excluded.

The motivations for the proposed transformation are varied: i) the transformed data doesn’t lose interpretability since the relation between the new features and the corresponding gene expression values is known; ii) the data stays in tabular form, allowing the use of classical ML models; iii) the new features encode relevant relations between the genes, simplifying the prediction task for classifiers; and iv) the identified putative regulatory modules can be used in a descriptive analysis of the data to strengthen the understanding of the condition under study.

### 2.4. Hyperparemterization

The proposed transformation has multiple hyperparameters that can be adjusted to best suit the data in question. These can be divided into two groups, the parameters of the biclustering algorithm and the parameters specific to the subsequent transformation.

The biclustering parameters subjected to optimization are: *number of iterations*, the number of repetitions of the mining process, with each new iteration masking the previously found biclusters to force the discovery of less trivial ones; *minimum lift*, a threshold that every bicluster must reach to be selected; *coherence strength*, the number of intervals used for the discretization of the real-valued data; and *minimum number of biclusters*, the number of biclusters per iteration at which the search stops.

The remaining hyperparameters are used directly in the transformation, comprising the filtering of the found patterns and the distance calculus, as previously explained.

Regarding biclustering, its parametrization was fixed according to sensitivity analysis over multiple cancers and omics layers. In a biological system, the underlying molecular mechanisms are complex, a result of the many interactions between a lot of molecules, be it transcripts, proteins, or others. Consequently, it is desirable to capture patterns across the whole dataset, being careful not to extract patterns focused only on a part of the data. Still, these patterns should be discriminative of the target variable, which requires a balance between the breadth of the search and the identification of patterns that are indeed relevant. Finally, we also aim for a significant compression of the existent information, in the form of dimensionality reduction. To achieve all this, the bicluster parameters are fixed as follows: a relatively high number of iterations (10) to force a broad search, a minimum lift of 1.2 to ensure the discriminative power of the patterns, a low minimum number of biclusters (10) to help reduce dimensionality and maintain the search feasible, and a heightened coherence strength (11) to ensure discrimination of the different values.

### 2.5. Implementation details

The proposed transformation is developed in Python 3.10, and the ML models used to assess classification performance are implemented in the scikit-learn package (Pedregosa et al.). The biclustering algorithm, BicPAMS (Henriques et al., 2017), is implemented in Java and the Differential Gene Expression (DEG) analysis is performed through the R package edgeR (Robinson et al., 2010). The code used to obtain the presented results is available in GitHub (github.com/Andrempp/Pattern-Centric-Transformation) and was run on a machine with an Intel Core i7-11700K 8-Core 3.6GHz and 32 GB of RAM.

## 3. Results

### 3.1. Evaluation and Metrics

To evaluate the impact of the proposed transformation on classification performance we train and test three ML classifiers: a Logistic Regression (LR), a Support Vector Machine (SVM), and a Random Forest (RF). This set of classifiers enables a broader understanding of the impacts on performance across different types of models, namely, linear (LR), kernel-based (SVM), and ensemble (RF). The evaluation is performed with a cross-validation setting with 10 folds, assuring that the patterns used for the transformation are discovered only on the training data.

The classifiers are evaluated through the following metrics: accuracy, to provide an overall indicator of performance; and recall, precision, and F1-score, to obtain a balanced view of how well the models identify the positive class (*Vital Status*).

To provide a reference of comparison for the results obtained in the transformed data, we also display the results obtained in the non-transformed dataset that maintains the transcripts/proteins as features. These results are presented in two settings. First, the results obtained in the miRNA and protein layers, where the genecentric dataset goes through the preprocessing steps previously explained. Second, the results obtained in the mRNA layer, where due to its high dimensionality the gene-centric dataset is further reduced through Differential Expression Gene (DEG) analysis with a *p*-value of 0.05 adjusted with Benjamini-Hochberg procedure multiple testing correction (Benjamini and Hochberg, 1995).

### 3.2. Prediction Results

The prediction results for Low Grade Glioma (LGG) can be seen in Figure 2 for miRNA and Figure 3 for protein.

**Fig. 2:**
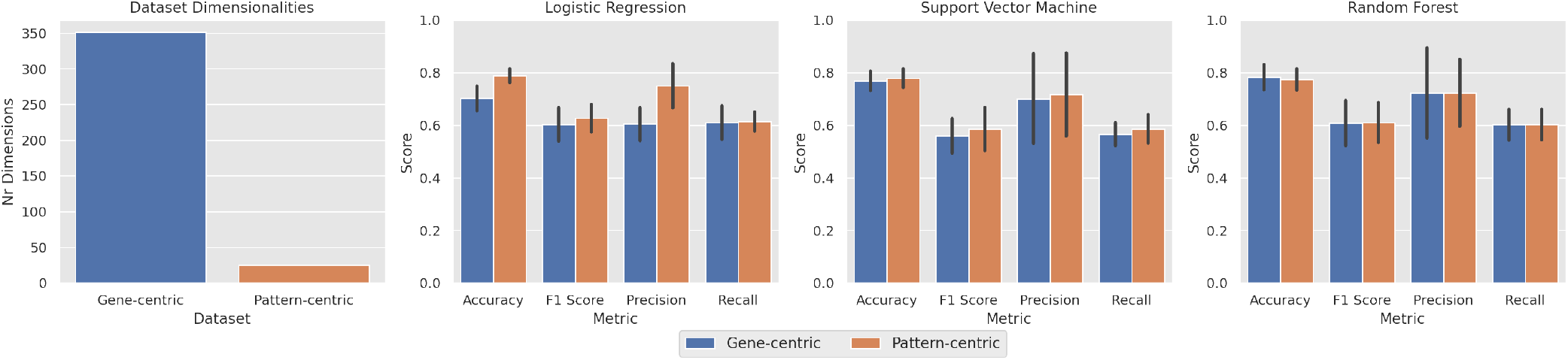
Comparison of dimensionality (left plot) and classification performance between gene-centric and pattern-centric data in the miRNA layer of TCGA-LGG project.

**Fig. 3:**
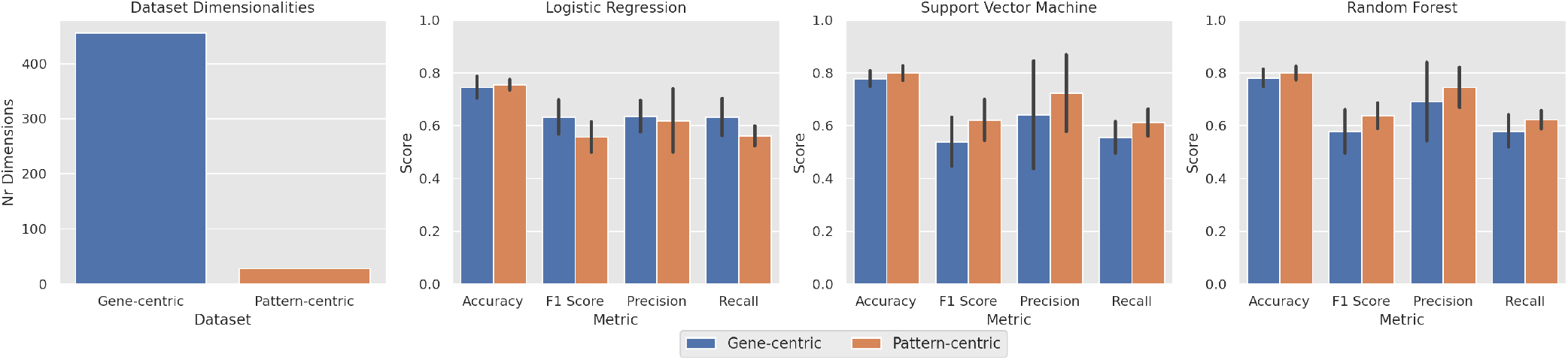
Comparison of dimensionality (left plot) and classification performance between gene-centric and pattern-centric data in the protein layer of TCGA-LGG project.

In the miRNA layer, averaging over the 10 folds of crossvalidation, there are 351 features in the gene-centric dataset and 26 features in the pattern-centric dataset. This corresponds to a reduction in the dimensionality of 92.6% while improving the predictive performance across all metrics and classifiers except one (accuracy in Random Forest). In the protein layer, there is a reduction of 457 features to 29, a 93.7% dimensionality reduction while improving the predictive performance of SVMs and Random Forests across all metrics.

Advancing to the mRNA layer, the dataset has a total of 60664 features before processing, and 24027 features postprocessing. In order to obtain a better comparison of results, the genecentric data is further reduced through feature selection with

DEG analysis, selecting the features that have a *p*-value of 0.05 or lower. The pattern-centric data is obtained by applying the transformation to the original preprocessed data, as in the previous results. Figure 4 shows the obtained results.

**Fig. 4:**
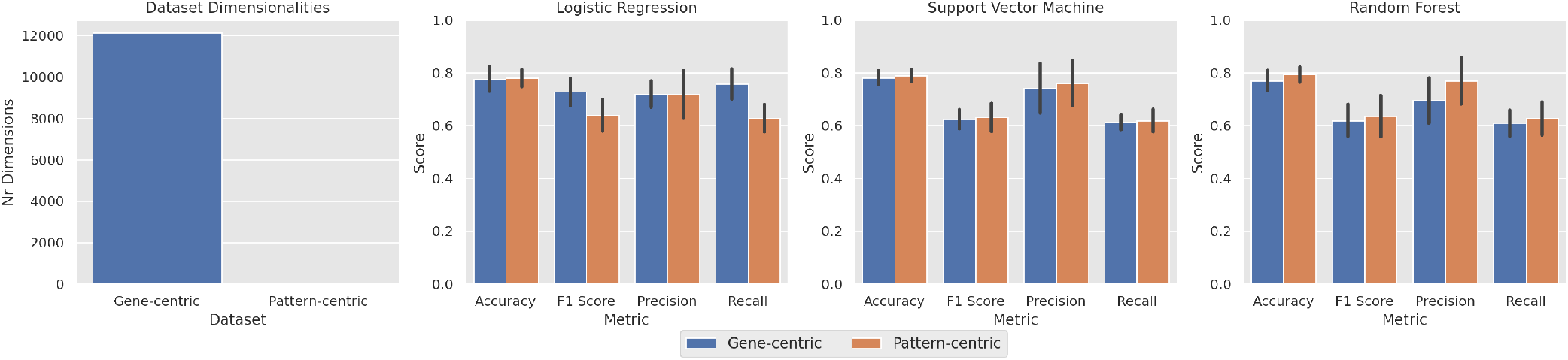
Comparison of dimensionality (left plot) and classification performance between gene-centric and pattern-centric data in the mRNA layer.

In the original dataset, the average dimensionality across folds after DEG analysis is 12140. The pattern-centric transformation reduces it to an average of 30 variables, corresponding to an increased reduction in the dimensionality of 99%. This is accompanied by an increase in performance in all metrics in the SVM and Random Forest classifiers. The Logistic Regression shows decreases in recall and F1-score.

In the TCGA-COAD project, increases in performance are observed, although less prominent. Furthermore, the reductions in dimensionality are above 80%. The complete prediction results can be consulted in Tables 2 and 3, and the corresponding dimensionality reductions are available in Table 4.

**Table 2.**
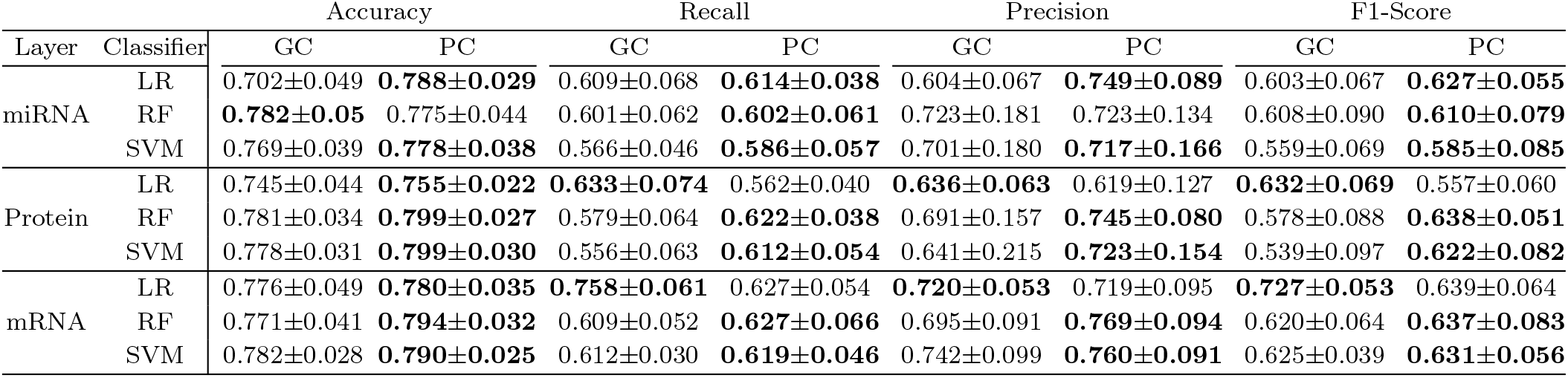
Comparison of classification results for different omics layers and classifiers between gene-centric (GC) and pattern-centric (PC) data in the TCGA-LGG project (best results in bold and ± are the standard deviations).

| Layer | Classifier | Accuracy |  | Recall |  | Precision |  | F1-Score |  |
| --- | --- | --- | --- | --- | --- | --- | --- | --- | --- |
|  |  | GC | PC | GC | PC | GC | PC | GC | PC |
| miRNA | LR | 0.702±0.049 | **0.788±0.029** | 0.609±0.068 | **0.614±0.038** | 0.604±0.067 | **0.749±0.089** | 0.603±0.067 | **0.627±0.055** |
| | RF | **0.782±0.05** | 0.775±0.044 | 0.601±0.062 | **0.602±0.061** | 0.723±0.181 | 0.723±0.134 | 0.608±0.090 | **0.610±0.079** |
| | SVM | 0.769±0.039 | **0.778±0.038** | 0.566±0.046 | **0.586±0.057** | 0.701±0.180 | **0.717±0.166** | 0.559±0.069 | **0.585±0.085** |
| Protein | LR | 0.745±0.044 | **0.755±0.022** | **0.633±0.074** | 0.562±0.040 | **0.636±0.063** | 0.619±0.127 | **0.632±0.069** | 0.557±0.060 |
| | RF | 0.781±0.034 | **0.799±0.027** | 0.579±0.064 | **0.622±0.038** | 0.691±0.157 | **0.745±0.080** | 0.578±0.088 | **0.638±0.051** |
| | SVM | 0.778±0.031 | **0.799±0.030** | 0.556±0.063 | **0.612±0.054** | 0.641±0.215 | **0.723±0.154** | 0.539±0.097 | **0.622±0.082** |
| mRNA | LR | 0.776±0.049 | **0.780±0.035** | **0.758±0.061** | 0.627±0.054 | **0.720±0.053** | 0.719±0.095 | **0.727±0.053** | 0.639±0.064 |
| | RF | 0.771±0.041 | **0.794±0.032** | 0.609±0.052 | **0.627±0.066** | 0.695±0.091 | **0.769±0.094** | 0.620±0.064 | **0.637±0.083** |
| | SVM | 0.782±0.028 | **0.790±0.025** | 0.612±0.030 | **0.619±0.046** | 0.742±0.099 | **0.760±0.091** | 0.625±0.039 | **0.631±0.056** |

**Table 3.**
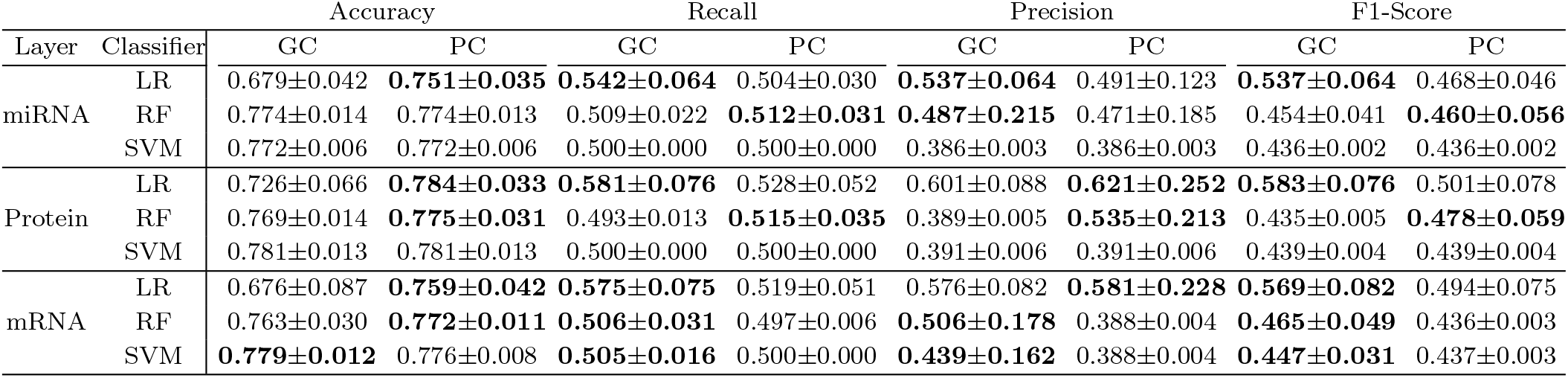
Comparison of classification results for different omics layers and classifiers between gene-centric (GC) and pattern-centric (PC) data in the TCGA-COAD project (best results in bold and ± are the standard deviations).

| Layer | Classifier | Accuracy |  | Recall |  | Precision |  | F1-Score |  |
| --- | --- | --- | --- | --- | --- | --- | --- | --- | --- |
|  |  | GC | PC | GC | PC | GC | PC | GC | PC |
| miRNA | LR | 0.679±0.042 | **0.751±0.035** | **0.542±0.064** | 0.504±0.030 | **0.537±0.064** | 0.491±0.123 | **0.537±0.064** | 0.468±0.046 |
| | RF | 0.774±0.014 | 0.774±0.013 | 0.509±0.022 | **0.512±0.031** | **0.487±0.215** | 0.471±0.185 | 0.454±0.041 | **0.460±0.056** |
| | SVM | 0.772±0.006 | 0.772±0.006 | 0.500±0.000 | 0.500±0.000 | 0.386±0.003 | 0.386±0.003 | 0.436±0.002 | 0.436±0.002 |
| Protein | LR | 0.726±0.066 | **0.784±0.033** | **0.581±0.076** | 0.528±0.052 | 0.601±0.088 | **0.621±0.252** | **0.583±0.076** | 0.501±0.078 |
| | RF | 0.769±0.014 | **0.775±0.031** | 0.493±0.013 | **0.515±0.035** | 0.389±0.005 | **0.535±0.213** | 0.435±0.005 | **0.478±0.059** |
| | SVM | 0.781±0.013 | 0.781±0.013 | 0.500±0.000 | 0.500±0.000 | 0.391±0.006 | 0.391±0.006 | 0.439±0.004 | 0.439±0.004 |
| mRNA | LR | 0.676±0.087 | **0.759±0.042** | **0.575±0.075** | 0.519±0.051 | 0.576±0.082 | **0.581±0.228** | **0.569±0.082** | 0.494±0.075 |
| | RF | 0.763±0.030 | **0.772±0.011** | **0.506±0.031** | 0.497±0.006 | **0.506±0.178** | 0.388±0.004 | **0.465±0.049** | 0.436±0.003 |
| | SVM | **0.779±0.012** | 0.776±0.008 | **0.505±0.016** | 0.500±0.000 | **0.439±0.162** | 0.388±0.004 | **0.447±0.031** | 0.437±0.003 |

**Table 4.** Dimensionalities before transformation (BT) and after transformation (AT) across different layers and projects.

| Project | miRNA |  |  | Protein |  |  | mRNA |  |  |
| --- | --- | --- | --- | --- | --- | --- | --- | --- | --- |
|  | BT | AT | Reduction | BT | AT | Reduction | BT | AT | Reduction |
| TCGA-LGG | 351 | 26 | 92.6% | 457 | 29 | 93.7% | 12140 | 30 | 99.7% |
| TCGA-COAD | 292 | 46 | 84.2% | 456 | 32 | 93.0% | 3720 | 41 | 98.9% |

### 3.3. Dimensionality Reduction

To better understand the capabilities of the proposed transformation for reducing dimensionality while preserving relevant information, we now consider the mRNA layer of the LGG project. Using the same evaluation setting as before, we compare the differences in performance in relation to a continuous decrease in the number of variables. We present the study of three techniques: DEG analysis, Principal Component Analysis (PCA), and the proposed pattern-centric transformation. For the DEG analysis, the variables are ordered according to their Benjamini-Hochberg adjusted *p*-value and the top *n* variables selected. This dimensionality reduction maintains the data in the same type of space, with the mRNA transcripts as the variables. The PCA transformation identifies the *n* top principal components, extracting information about the variability of the data and using it to transform it. This results in an alteration of the variable space to one that hinders human interpretation. Regarding the pattern-centric transformation, it is applied as previously explained, but it further filters the found patterns according to their lift, enabling a further reduction of dimensionality. This process transforms the variable space into a different one, where each variable represents the interactions between the original variables, whilst maintaining the possibility of human interpretation. The obtained results can be seen in Figure 5.

**Fig. 5:**
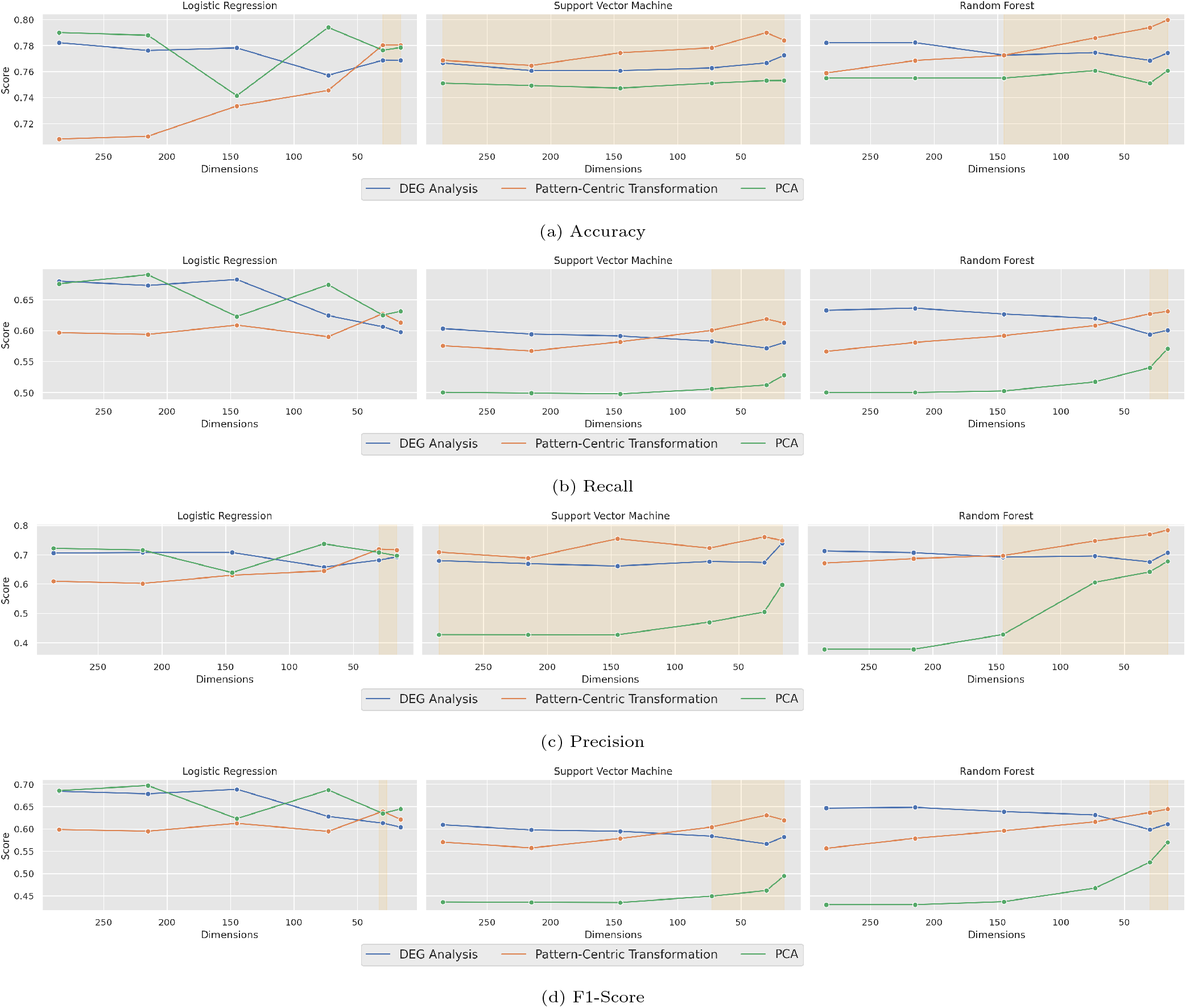
Classification performance of distinct dimensionality reduction methods across different dimensionalities (better performance by Pattern-Centric Transformation highlighted in orange).

The results show that although the pattern-centric transformation doesn’t start as the best-performing technique, as the dimensionality is reduced, its performance steadily increases, making it the best performer at the lowest dimensionalities.

### 3.4. Feature Space

In addition to the impact on performance, it is also useful to analyze how the proposed transformation affects other characteristics of the feature space, namely, class separability.

Both the gene- and pattern-centric data are reduced to two dimensions through the UMAP (McInnes et al., 2020) technique and their data points are plotted using color to differentiate the two classes (*Alive* and *Deceased*), as seen in Figure 6. Both plots indicate an inherently difficult classification task since there is no clear separation between the two classes. Still, it is visible that the transformation causes a significant alteration in the way data is distributed.

**Fig. 6:**
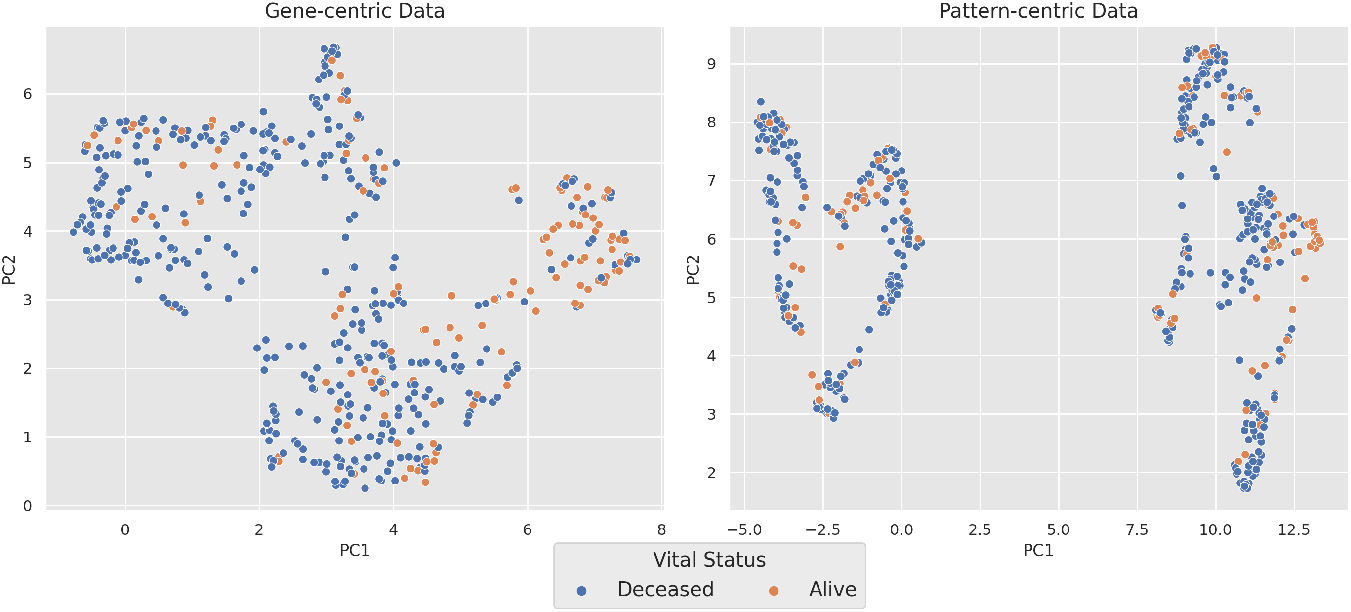
Class separability in two dimensions for the gene-centric (left) and pattern-centric (right) data.

Complementary to the study of the data distribution as a whole, the distribution of individual variables is also analyzed.

We compare the distributions conditioned on the target variable of the most discriminative features, both in the gene- and pattern-centric datasets. The five top discriminative features for each dataset are selected through their Mutual Information (MI). Figure 7 shows a Kernel Density Estimate (KDE) plot for the conditioned distributions of each of these variables, with the genecentric features on top and the pattern-centric features on the bottom. The overlap of the conditioned distributions in the genecentric dataset indicates, once again, an innate difficulty in this prediction task. However, the pattern-centric features show a clear improvement in the separability of the two distributions, demonstrating that the new feature space yields higher guarantees of discriminative power.

**Fig. 7:**
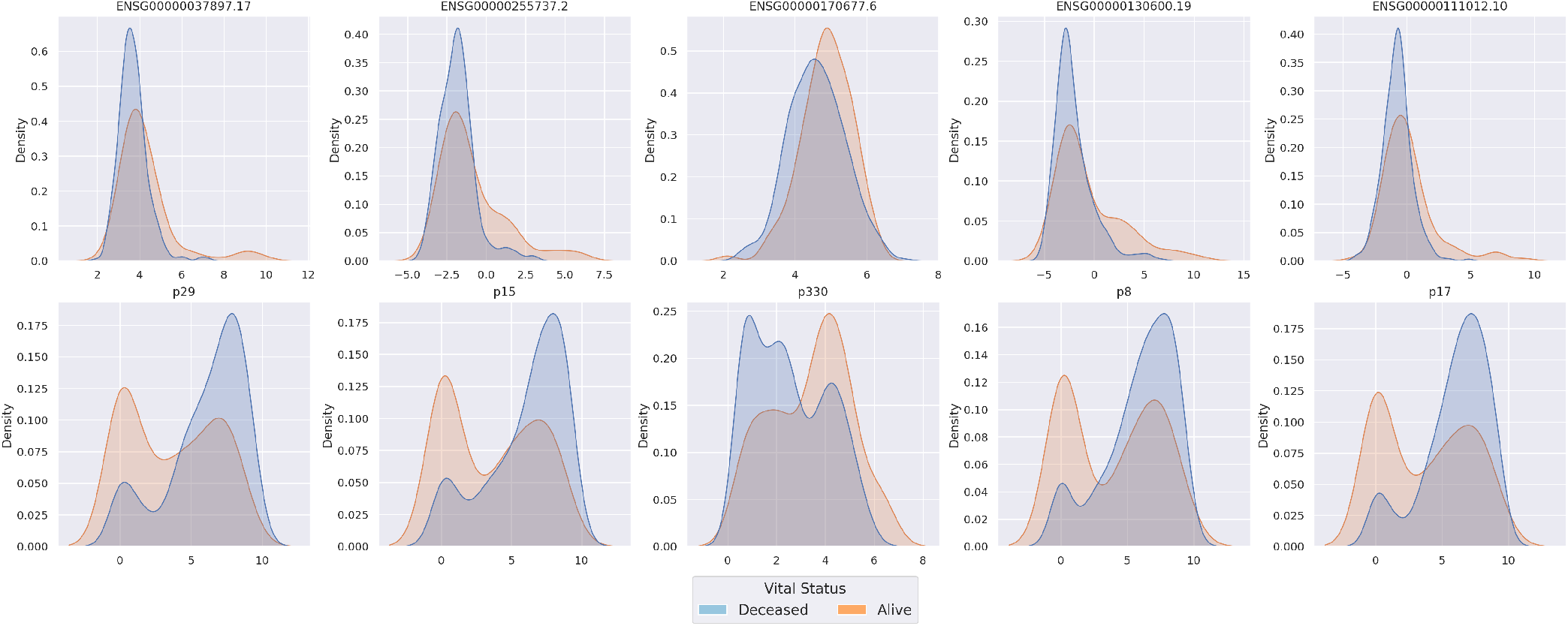
Normalized conditional distribution of the most discriminative variables in gene-centric (top) and pattern-centric (bottom) data.

The same analysis for the data from the project COAD can be found in Supplementary Figures S1 and S2.

## 4. Discussion

Omics data contains fundamental information about regulatory biological mechanisms, however, capturing the inherent complexity of molecular interplay in the various pathways is a challenging task. This complicates the use of omics data in clinical settings since it is important to maintain some level of interpretability of both the data and the obtained results. We have proposed here a transformation based on the mining of putative regulatory patterns with discriminative guarantees to create a new feature space with unique properties of interest.

Through the extraction of relevant relations from the data, this transformation can improve prediction capability, not only maintaining the interpretability according to the original variables but also increasing it. The underlying hypothesis is that such a transformation facilitates both predictive learning and the interpretation of biological mechanisms by humans.

The present results confirm that the transformation can indeed result in increased classification performance and additionally serve as an efficient feature extraction and dimensionality reduction technique.

The prediction results (Figures 2, 3 and 4 and Tables 2 and 3) show an improvement in performance across different layers and cancer types. In addition, we can further see a considerable compression of the information scattered throughout thousands of variables into the new features (Table 4), reaching in some cases, a reduction in dimensionality of 99%.

The impact of the data compression on classification performance can be further seen in Figure 5, where is shown that the transformation actually performs better with an increasingly reduced dimensionality. This can be explained by the way the biclustering algorithm works. Biclustering performs a defined number of iterations of its search, returning all patterns found above a defined lift, some considerably more discriminative than others. It is through the subsequent filtering step that we can obtain the patterns with higher hypothesized discriminative power whilst removing patterns that mainly contributed with noise, resulting in increased performance with lower dimensionalities.

We further show that this transformation creates variables that can more easily distinguish between the conditions under study, contrary to the original features that present highly overlapping conditional distributions (Figure 7). However, the discriminative power of the highest discriminative patterns is diluted when considering all patterns (Figure 6). Still, it is noted that the transformation has a strong impact on the distribution of the data points.

In addition to the listed benefits in prediction capability and dimensionality reduction, the proposed transformation can further be useful for a descriptive analysis of the found patterns, possibly leading to a deeper understanding of the conditions under study. As an example, we look at the pattern found to be the most discriminative in the mRNA layer of the LGG project, denominated as *p29* and whose conditional distribution is shown in Figure 7. This pattern is found in 52 out of the total 534 samples, corresponds to high expression levels of genes *EMP3, CHI3L1*, and *SH2D4A*, and presents a lift of 2.51. Functional enrichment analysis performed through the Enrichr tool (Chen et al., 2013) shows a relation between this gene set and a pathway for Glioblastoma according to the Elsevier Pathway Collection.

Furthermore, patterns *p15* (genes *TAGLN2, TIMP1*, and *EMP3*) and *p330* (genes *HOXA11-AS, HOXA7*, and *NTS*), also present in Figure 7, show relations to pathways concerning proteins involved in Glioblastoma and telomerase reverse transcriptase (TERT) activation in cancer, respectively.

The results here presented open the door to pattern-centric transformations with patterns obtained through methods other than biclustering since as long as the found patterns are significantly discriminative, some benefits will arguably hold. Even in the context of biclustering, there are other alternatives that should be tested, namely, the use of non-constant patterns. Order-preserving biclusters, for example, would capture just the relations of order between genes, adjusting for different base levels of expression between individuals, while coherent-valued biclusters would further enforce that the found patterns follow coherent variations across features. The proposed methodology can be further extended in multiple ways, namely: by changing the method to assess the level of expression of a given pattern in a sample or by filtering the found patterns by other characteristics such as statistical significance and putative biological relevance.

## Author contributions statement

Conceptualization: all authors; Methodology: A.P., R.H.; Formal analysis: A.P.; Writing - original draft: A.P.; Funding Acquisition: R.H., R.S.C.; Writing - review and editing: all authors; Supervision: R.S.C., R.H. All authors have read and agreed to the published version of the manuscript.

## Data availability

The data underlying this article are available in GDC Data Portal at portal.gdc.cancer.gov, and can be accessed with the project codes TCGA-LGG and TCGA-COAD. The processed data is available at **github**.

## Footnotes

1 https://portal.gdc.cancer.gov/projects/TCGA-LGG

2 https://portal.gdc.cancer.gov/projects/TCGA-COAD

## References

S. Acharya, S. Saha, and N. Nikhil. Unsupervised gene selection using biological knowledge : Application in sample clustering. BMC Bioinformatics, 18(1):513, Nov. 2017. ISSN 1471-2105. doi: 10.1186/s12859-017-1933-0.

Y. Benjamini and Y. Hochberg. Controlling the false discovery rate: A practical and powerful approach to multiple testing. Journal of the Royal Statistical Society. Series B. Methodological, 57(1):289–300, 1995. ISSN 0035-9246.

B. Chandra and M. Gupta. An efficient statistical feature selection approach for classification of gene expression data. Journal of Biomedical Informatics, 44(4):529–535, Aug. 2011. ISSN 1532-0464. doi: 10.1016/j.jbi.2011.01.001.

E. Y. Chen, C. M. Tan, Y. Kou, Q. Duan, Z. Wang, G. V. Meirelles, N. R. Clark, and A. Ma’ayan. Enrichr: Interactive and collaborative HTML5 gene list enrichment analysis tool. BMC bioinformatics, 14:128, Apr. 2013. ISSN 1471-2105. doi: 10.1186/1471-2105-14-128.

Y. Chen, A. T. L. Lun, and G. K. Smyth. From reads to genes to pathways: Differential expression analysis of RNA-Seq experiments using Rsubread and the edgeR quasi-likelihood pipeline. F1000Research, 5:1438, Aug. 2016. ISSN 2046-1402. doi: 10.12688/f1000research.8987.2.

R. Clarke, H. W. Ressom, A. Wang, J. Xuan, M. C. Liu, E. A. Gehan, and Y. Wang. The properties of high-dimensional data spaces: Implications for exploring gene and protein expression data. Nature reviews. Cancer, 8(1):37–49, Jan. 2008. ISSN 1474-175X. doi: 10.1038/nrc2294.

A. Colaprico, T. C. Silva, C. Olsen, L. Garofano, C. Cava, D. Garolini, T. S. Sabedot, T. M. Malta, S. M. Pagnotta, I. Castiglioni, M. Ceccarelli, G. Bontempi, and H. Noushmehr. TCGAbiolinks: An R/Bioconductor package for integrative analysis of TCGA data. Nucleic Acids Research, 44(8):e71, May 2016. ISSN 0305-1048. doi: 10.1093/nar/gkv1507.

C. Ding and H. Peng. Minimum redundancy feature selection from microarray gene expression data. Journal of Bioinformatics and Computational Biology, 03(02):185–205, Apr. 2005. ISSN 0219-7200. doi: 10.1142/S0219720005001004.

A.-L. Doebley, M. Ko, H. Liao, A. E. Cruikshank, K. Santos, C. Kikawa, J. B. Hiatt, R. D. Patton, N. De Sarkar, K. A. Collier, A. C. H. Hoge, K. Chen, A. Zimmer, Z. T. Weber, M. Adil, J. B. Reichel, P. Polak, V. A. Adalsteinsson, P. S. Nelson, D. MacPherson, H. A. Parsons, D. G. Stover, and G. Ha. A framework for clinical cancer subtyping from nucleosome profiling of cell-free DNA. Nature Communications, 13(1):7475, Dec. 2022. ISSN 2041-1723. doi: 10.1038/s41467-022-35076-w.

S. Fan, J. Tang, Q. Tian, and C. Wu. A robust fuzzy rule based integrative feature selection strategy for gene expression data in TCGA. BMC Medical Genomics, 12(1):14, Jan. 2019. ISSN 1755-8794. doi: 10.1186/s12920-018-0451-x.

J. Ferlay, M. Colombet, I. Soerjomataram, D. M. Parkin, M. Piñeros, A. Znaor, and F. Bray. Cancer statistics for the year 2020: An overview. International Journal of Cancer, 149 (4):778–789, 2021. ISSN 1097-0215. doi: 10.1002/ijc.33588.

O. Gevaert, F. D. Smet, D. Timmerman, Y. Moreau, and B. D. Moor. Predicting the prognosis of breast cancer by integrating clinical and microarray data with Bayesian networks. Bioinformatics, 22(14):e184–e190, July 2006. ISSN 1367-4803. doi: 10.1093/bioinformatics/btl230.

S. Ghosal, S. Das, Y. Pang, M. K. Gonzales, T.-T. Huynh, Y. Yang, D. Taieb, J. Crona, U. T. Shankavaram, and K. Pacak. Long intergenic noncoding RNA profiles of pheochromocytoma and paraganglioma: A novel prognostic biomarker. International Journal of Cancer, 146(8):2326–2335, 2020. ISSN 1097-0215. doi: 10.1002/ijc.32654.

R. Henriques and S. C. Madeira. BicPAM: Pattern-based biclustering for biomedical data analysis. Algorithms for Molecular Biology, 9(1):27, Dec. 2014. ISSN 1748-7188. doi: 10.1186/s13015-014-0027-z.

R. Henriques and S. C. Madeira. BSig: Evaluating the statistical significance of biclustering solutions. Data Mining and Knowledge Discovery, 32(1):124–161, Jan. 2018. ISSN 1573-756X. doi: 10.1007/s10618-017-0521-2.

R. Henriques and S. C. Madeira. FleBiC: Learning classifiers from high-dimensional biomedical data using discriminative biclusters with non-constant patterns. Pattern Recognition, 115: 107900, July 2021. ISSN 0031-3203. doi: 10.1016/j.patcog.2021.107900.

R. Henriques, C. Antunes, and S. C. Madeira. A structured view on pattern mining-based biclustering. Pattern Recognition, 48 (12):3941–3958, Dec. 2015. ISSN 0031-3203. doi: 10.1016/j.patcog.2015.06.018.

R. Henriques, F. L. Ferreira, and S. C. Madeira. BicPAMS: Software for biological data analysis with pattern-based biclustering. BMC Bioinformatics, 18(1):82, Feb. 2017. ISSN 1471-2105. doi: 10.1186/s12859-017-1493-3.

C.-T. Huang, C.-H. Hsieh, Y.-H. Chung, Y.-J. Oyang, H.-C. Huang, and H.-F. Juan. Perturbational Gene-Expression Signatures for Combinatorial Drug Discovery. iScience, 15:291–306, May 2019. ISSN 2589-0042. doi: 10.1016/j.isci.2019.04.039.

Y. Li, D. M. Umbach, J. M. Krahn, I. Shats, X. Li, and L. Li. Predicting tumor response to drugs based on gene-expression biomarkers of sensitivity learned from cancer cell lines. BMC Genomics, 22(1):272, Apr. 2021. ISSN 1471-2164. doi: 10.1186/s12864-021-07581-7.

H. Lu, J. Chen, K. Yan, Q. Jin, Y. Xue, and Z. Gao. A hybrid feature selection algorithm for gene expression data classification. Neurocomputing, 256:56–62, Sept. 2017. ISSN 0925-2312. doi: 10.1016/j.neucom.2016.07.080.

S. Madeira and A. Oliveira. Biclustering algorithms for biological data analysis: A survey. IEEE/ACM Transactions on Computational Biology and Bioinformatics, 1(1):24–45, Jan. 2004. ISSN 1557-9964. doi: 10.1109/TCBB.2004.2.

J. U. Marquardt, P. R. Galle, and A. Teufel. Molecular diagnosis and therapy of hepatocellular carcinoma (HCC): An emerging field for advanced technologies. Journal of Hepatology, 56(1): 267–275, Jan. 2012. ISSN 0168-8278. doi: 10.1016/j.jhep.2011.07.007.

L. McInnes, J. Healy, and J. Melville. UMAP: Uniform Manifold Approximation and Projection for Dimension Reduction, Sept. 2020.

P. I. Metselaar, L. Mendoza-Maldonado, A. Y. F. Li Yim, I. Abarkan, P. Henneman, A. A. te Velde, A. Schönhuth, J. A. Bosch, A. D. Kraneveld, and A. Lopez-Rincon. Recursive ensemble feature selection provides a robust mRNA expression signature for myalgic encephalomyelitis/chronic fatigue syndrome. Scientific Reports, 11(1):4541, Feb. 2021. ISSN 2045-2322. doi: 10.1038/s41598-021-83660-9.

J.-C. Nault, A. De Reyniés, A. Villanueva, J. Calderaro, S. Rebouissou, G. Couchy, T. Decaens, D. Franco, S. Imbeaud, F. Rousseau, D. Azoulay, J. Saric, J.-F. Blanc, C. Balabaud, P. Bioulac-Sage, A. Laurent, P. Laurent-Puig, J. M. Llovet, and J. Zucman-Rossi. A hepatocellular carcinoma 5-gene score associated with survival of patients after liver resection. Gastroenterology, 145(1):176–187, July 2013. ISSN 1528-0012. doi: 10.1053/j.gastro.2013.03.051.

B. Neary, J. Zhou, and P. Qiu. Identifying gene expression patterns associated with drug-specific survival in cancer patients. Scientific Reports, 11(1):5004, Mar. 2021. ISSN 2045-2322. doi: 10.1038/s41598-021-84211-y.

F. Pedregosa, G. Varoquaux, A. Gramfort, V. Michel, B. Thirion, O. Grisel, M. Blondel, P. Prettenhofer, R. Weiss, V. Dubourg, J. Vanderplas, A. Passos, and D. Cournapeau. Scikit-learn: Machine Learning in Python. MACHINE LEARNING IN PYTHON.

V. K. Raghu, X. Ge, P. K. Chrysanthis, and P. V. Benos. Integrated Theory-and Data-Driven Feature Selection in Gene Expression Data Analysis. In 2017 IEEE 33rd International Conference on Data Engineering (ICDE), pages 1525–1532, Apr. 2017. doi: 10.1109/ICDE.2017.223.

M. D. Robinson, D. J. McCarthy, and G. K. Smyth. edgeR: A Bioconductor package for differential expression analysis of digital gene expression data. Bioinformatics, 26(1):139–140, Jan. 2010. ISSN 1367-4803. doi: 10.1093/bioinformatics/btp616.

S. Roessler, H.-L. Jia, A. Budhu, M. Forgues, Q.-H. Ye, J.-S. Lee, S. S. Thorgeirsson, Z. Sun, Z.-Y. Tang, L.-X. Qin, and X. W. Wang. A Unique Metastasis Gene Signature Enables Prediction of Tumor Relapse in Early Stage Hepatocellular Carcinoma Patients. Cancer research, 70(24):10202–10212, Dec. 2010. ISSN 008-5472. doi: 10.1158/0008-5472.CAN-10-2607.

F. Rohart, B. Gautier, A. Singh, and K.-A. L. Cao. mixOmics: An R package for ‘omics feature selection and multiple data integration. PLOS Computational Biology, 13(11):e1005752, Nov. 2017. ISSN 1553-7358. doi: 10.1371/journal.pcbi.1005752.

Y. Saeys, I. Inza, and P. Larrañaga. A review of feature selection techniques in bioinformatics. Bioinformatics, 23(19):2507–2517, Oct. 2007. ISSN 1367-4803. doi: 10.1093/bioinformatics/btm344.

P. Sun, Y. Wu, C. Yin, H. Jiang, Y. Xu, and H. Sun. Molecular Subtyping of Cancer Based on Distinguishing Co-Expression Modules and Machine Learning. Frontiers in Genetics, 13, 2022. ISSN 1664-8021.

A. L. Swan, A. Mobasheri, D. Allaway, S. Liddell, and J. Bacardit. Application of Machine Learning to Proteomics Data: Classification and Biomarker Identification in Postgenomics Biology. OMICS: A Journal of Integrative Biology, 17(12): 595–610, Dec. 2013. doi: 10.1089/omi.2013.0017.

J. N. Weinstein, E. A. Collisson, G. B. Mills, K. M. Shaw, B. A. Ozenberger, K. Ellrott, I. Shmulevich, C. Sander, and J. M. Stuart. The Cancer Genome Atlas Pan-Cancer Analysis Project. Nature genetics, 45(10):1113–1120, Oct. 2013. ISSN 1061-4036. doi: 10.1038/ng.2764.

K. Y. Yeung and R. E. Bumgarner. Multiclass classification of microarray data with repeated measurements: Application to cancer. Genome Biology, 4(12):R83, Nov. 2003. ISSN 1474-760X. doi: 10.1186/gb-2003-4-12-r83.

M. Yousef, B. Bakir-Gungor, A. Jabeer, G. Goy, R. Qureshi, and L. C. Showe. Recursive Cluster Elimination based Rank Function (SVM-RCE-R) implemented in KNIME, Jan. 2021a.

M. Yousef, A. Kumar, and B. Bakir-Gungor. Application of Biological Domain Knowledge Based Feature Selection on Gene Expression Data. Entropy, 23(1):2, Jan. 2021b. ISSN 1099-4300. doi: 10.3390/e23010002.

T. Zhang, Y. Hu, W. Jiang, L. Fang, X. Guan, J. Chen, J. Zhang, C. A. Saski, B. E. Scheffler, D. M. Stelly, A. M. Hulse-Kemp, Q. Wan, B. Liu, C. Liu, S. Wang, M. Pan, Y. Wang, D. Wang, W. Ye, L. Chang, W. Zhang, Q. Song, R. C. Kirkbride, X. Chen, E. Dennis, D. J. Llewellyn, D. G. Peterson, P. Thaxton, D. C. Jones, Q. Wang, X. Xu, H. Zhang, H. Wu, L. Zhou, G. Mei, S. Chen, Y. Tian, D. Xiang, X. Li, J. Ding, Q. Zuo, L. Tao, Y. Liu, J. Li, Y. Lin, Y. Hui, Z. Cao, C. Cai, X. Zhu, Z. Jiang, B. Zhou, W. Guo, R. Li, and Z. J. Chen. Sequencing of allotetraploid cotton (Gossypium hirsutum L. acc. TM-1) provides a resource for fiber improvement. Nature Biotechnology, 33(5):531–537, May 2015. ISSN 1546-1696. doi: 10.1038/nbt.3207.

Y. Zhang, J. Shi, X. Liu, L. Feng, Z. Gong, P. Koppula, K. Sirohi, X. Li, Y. Wei, H. Lee, L. Zhuang, G. Chen, Z.-D. Xiao, M.-C. Hung, J. Chen, P. Huang, W. Li, and B. Gan. BAP1 links metabolic regulation of ferroptosis to tumour suppression. Nature Cell Biology, 20(10):1181–1192, Oct. 2018. ISSN 1476-4679. doi: 10.1038/s41556-018-0178-0.

L. Zhao, V. H. F. Lee, M. K. Ng, H. Yan, and M. F. Bijlsma. Molecular subtyping of cancer: Current status and moving toward clinical applications. Briefings in Bioinformatics, 20(2): 572–584, Mar. 2019. ISSN 1477-4054. doi: 10.1093/bib/bby026.

X. Zhao, Y. Yang, B.-F. Sun, Y. Shi, X. Yang, W. Xiao, Y.-J. Hao, X.-L. Ping, Y.-S. Chen, W.-J. Wang, K.-X. Jin, X. Wang, C.-M. Huang, Y. Fu, X.-M. Ge, S.-H. Song, H. S. Jeong, H. Yanagisawa, Y. Niu, G.-F. Jia, W. Wu, W.-M. Tong Okamoto, C. He, J. M. R. Danielsen, X.-J. Wang, and Y.-G. Yang. FTO-dependent demethylation of N6-methyladenosine regulates mRNA splicing and is required for adipogenesis. Cell Research, 24(12):1403–1419, Dec. 2014. ISSN 1748-7838. doi: 10.1038/cr.2014.151.

X. Zhao, J. Dou, J. Cao, Y. Wang, Q. Gao, Q. Zeng, W. Liu, Liu, Z. Cui, L. Teng, J. Zhang, and C. Zhao. Uncovering the potential differentially expressed miRNAs as diagnostic biomarkers for hepatocellular carcinoma based on machine learning in The Cancer Genome Atlas database. Oncology Reports, 43(6):1771–1784, June 2020. ISSN 1021-335X. doi: 10.3892/or.2020.7551.

