## Supplemental Materials - Figures S1-2 for "Pattern-centric transformation of omics-data sources grounded on multi-wise gene associations aids predictive tasks in TCGA while ensuring interpretability"

### 1 Supplementary Figures

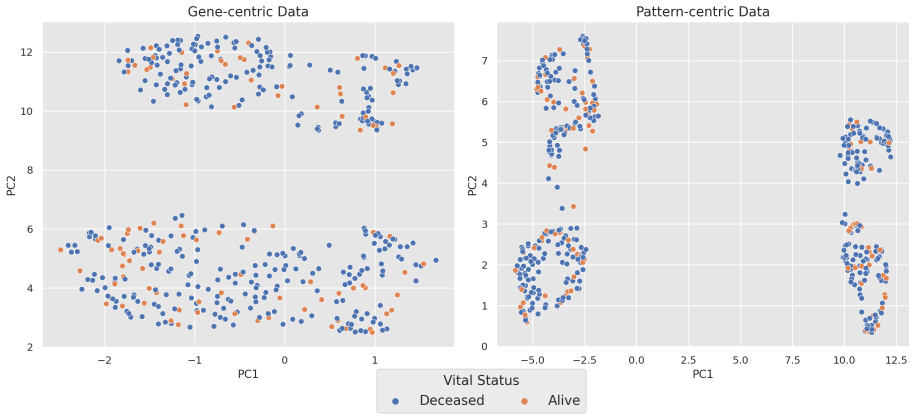


Figure S1: Class separability in two dimensions for the gene-centric (left) and pattern-centric (right) data in the TCGA-COAD project.

‘

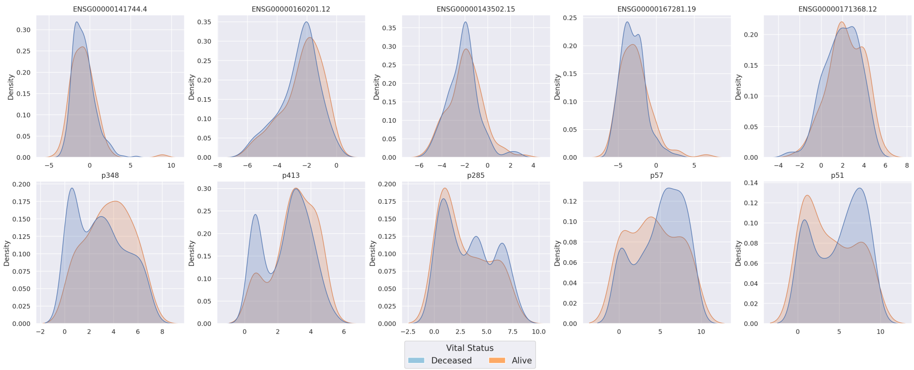


Figure S2: Normalized conditional distribution of the most discriminative variables in gene-centric (top) and pattern-centric (bottom) data in the TCGA-COAD project.
